# Automatic identification of drug-induced liver injury literature using natural language processing and machine learning methods

**DOI:** 10.1101/2022.08.10.503489

**Authors:** Jung Hun Oh, Allen Tannenbaum, Joseph O. Deasy

**Affiliations:** Department of Medical Physics, Memorial Sloan Kettering Cancer Center, New York, NY, USA; Departments of Computer Science and Applied Mathematics & Statistics, Stony Brook University, New York, NY, USA

## Abstract

Drug-induced liver injury (DILI) is an adverse hepatic drug reaction that can potentially lead to life-threatening liver failure. Previously published work in the scientific literature on DILI has provided valuable insights for the understanding of hepatotoxicity as well as drug development. However, the manual search of scientific literature in PubMed is laborious. Natural language processing (NLP) techniques have been developed to decipher and understand the meaning of human language by extracting useful information from unstructured text data. In particular, NLP along with artificial intelligence (AI) / machine learning (ML) techniques may allow automatic processing of the DILI literature, but useful methods are yet to be demonstrated. To address this challenge, we have developed an integrated NLP/ML classification model to identify DILI-related literature using only paper titles and abstracts. We used 14,203 publications provided by the Critical Assessment of Massive Data Analysis (CAMDA) challenge, employing word vectorization techniques in NLP coupled with machine learning methods. Classification modeling was performed using 2/3 of the data for training and the remainder for testing in internal validation. The best performance was achieved using a linear support vector machine (SVM) model that combined vectors derived from *term frequency-inverse document frequency (TF-IDF)* and *Word2Vec*, achieving an accuracy of 95.0% and an F1-score of 95.0%. The final SVM model built using all 14,203 publications was tested on independent datasets, resulting in accuracies of 92.5%, 96.3%, and 98.3%, and F1-scores of 93.5%, 86.1%, and 75.6% for three test sets (T1-T3). The SVM model was tested on four external validation sets (V1-V4), resulting in accuracies of 92.0%, 96.2%, 98.3%, and 93.1%, and F1-scores of 92.4%, 82.9%, 75.0%, and 93.3%.

## 1. Materials and Methods

### 1.1 Data

The Critical Assessment of Massive Data Analysis (CAMDA) 2022 in collaboration with the Intelligent Systems for Molecular Biology (ISMB) hosted the Literature AI for Drug Induced Liver Injury (DILI) challenge [1]. A curated dataset, consisting of 277,016 DILI annotated papers, was downloaded from the CAMDA website. All the papers were labeled as DILI-related (referred to as positive samples) or irrelevant to DILI (referred to as negative samples) by a panel of DILI experts. For the CAMDA challenge, the labels for 7,177 DILI-related papers and 7,026 DILI-unrelated papers were released while the labels for the remaining papers (N=262,813) were masked for model assessment and were split into three test sets and four validation sets. In this study, 14,203 papers with positive or negative labels were used for model building. For each paper, the title and abstract were provided and concatenated to be used in modeling.

### 1.2 Word vectorization

To extract text features from the unstructured literature, we employed two word vectorization techniques: word embedding with Word2Vec implemented in Gensim Python library [2] and Bag-of-Words (BoW) with term frequency-inverse document frequency (TF-IDF) weighting [3]. Before vectorization, text data were preprocessed by lowercasing, removing punctuations, special characters, white spaces, and standard stop-words followed by lemmatization, employing spaCy Python library [4]. The Word2Vec model was trained using the skip-gram method with a window size of 5 words and a 200-dimensional output vector for each word. The resulting output vectors for all the words in a given paper were averaged to create a single 200-dimensional vector to be used in modeling.

### 1.3 Modeling

Several machine learning methods, including support vector machine (SVM), logistic regression, and random forest were then tested on the transformed numerical features obtained from the Word2Vec and TF-IDF models. In addition, the Transformer-based BERT language model was tested for benchmark comparison [5]. For internal validation, in the machine learning modeling, the data of 14,203 samples were randomly stratified into the training (66.7%) and test (33.3%) sets. For the BERT model, the data were stratified into the training (56.7%), validation (10%), and test (33.3%) sets. This modeling process was iterated 30 times, and the averaged accuracy, precision, recall, and F1-score assessed on the test set were reported. For external validation, a final model was built using all the 14,203 samples and was tested on seven independent datasets (in total, N=262,813).

## 2. Results

Figure 1 illustrates the top 10 most common words in DILI-related (top) and unrelated (bottom) publications, with “patient” being the most frequent word in both DILI-related and unrelated publications. Note that “liver” and drug relevant words including “mg”, “drug”, and “dose” were the frequently occurring words in DILI-related literature, but were not included in the top 10 most common words in literature unrelated to DILI.

**Figure 1.**
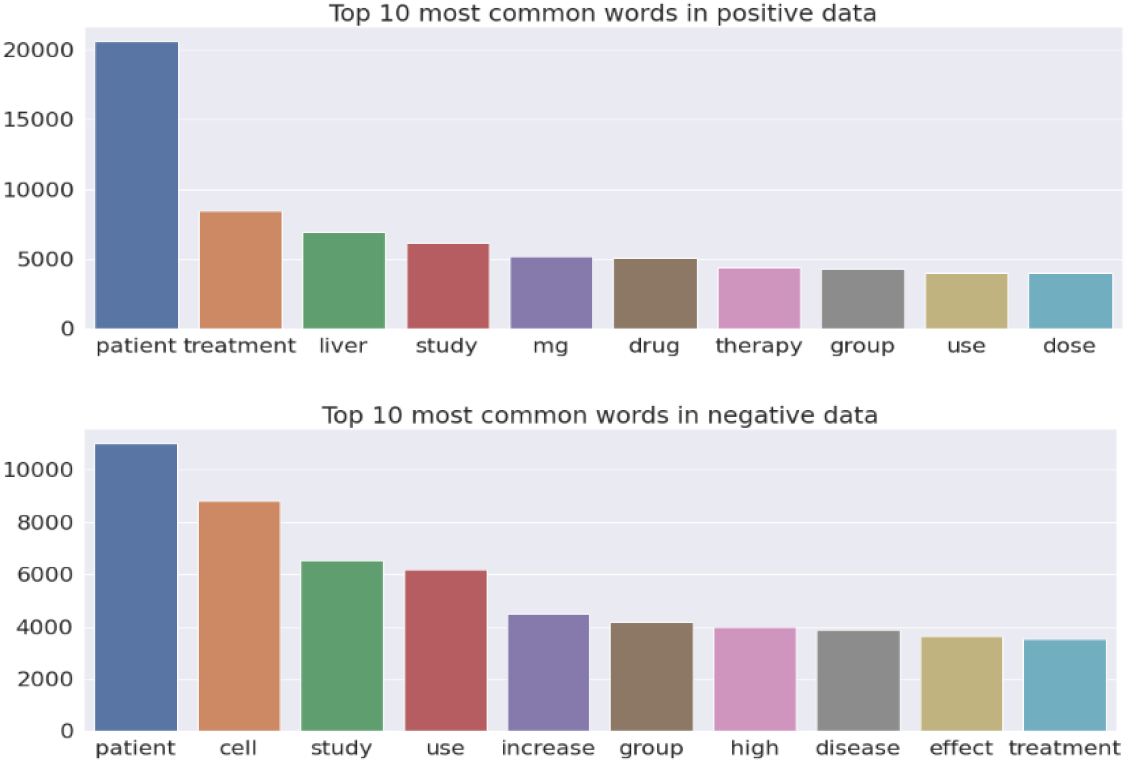
The top 10 most common words in (top) DILI-related and (bottom) unrelated literature.

Figure 2 illustrates the t-distributed stochastic neighbor embedding (t-SNE) visualization of the TF-IDF vectors obtained using the combined title and abstract vs only the title of each publication [6]. It is not surprising that the abstract has much more information than the title to distinguish DILI-related publications from the ones unrelated to DILI.

**Figure 2.**
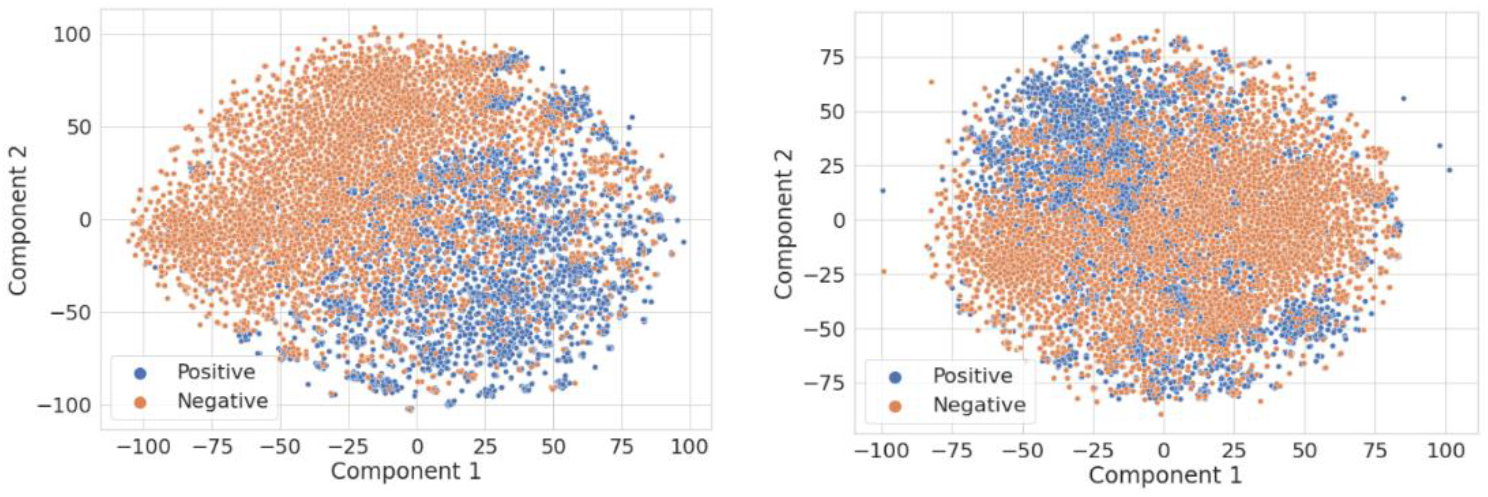
The t-SNE visualization of the TF-IDF vectors obtained using (left) the title and abstract and (right) only the title of each publication.

### 2.1 Internal validation

Figure 3 shows the accuracy of machine learning models on test data after 30 iterations, using vectors derived from the Word2Vec and TF-IDF models. A linear SVM model using vectors derived from the TF-IDF model achieved the best accuracy of 94.5%. Interestingly, a linear SVM using TF-IDF vectors derived from only the title of publications obtained an accuracy of 88.8%, whereas a linear SVM using TF-IDF vectors derived from only the abstract of publications obtained an accuracy of 94.3%, implying that the information available in the title of publications did not improve performance in classifying DILI-related literature. In addition, machine learning models were tested on the combined data of Word2Vec and TF-IDF vectors. A linear SVM model achieved the best performance with an accuracy of 95.0%, aprecision of 95.3%, arecall of 94.7%, and an F1-score of 95.0% (Table 1). Overall, this approach improved classification performance compared to the models on Word2Vec or TF-IDF vectors alone. A BERT model had worse performance in all metrics as compared to other machine learning models coupled with Word2Vec or TF-IDF techniques.

**Figure 3.**
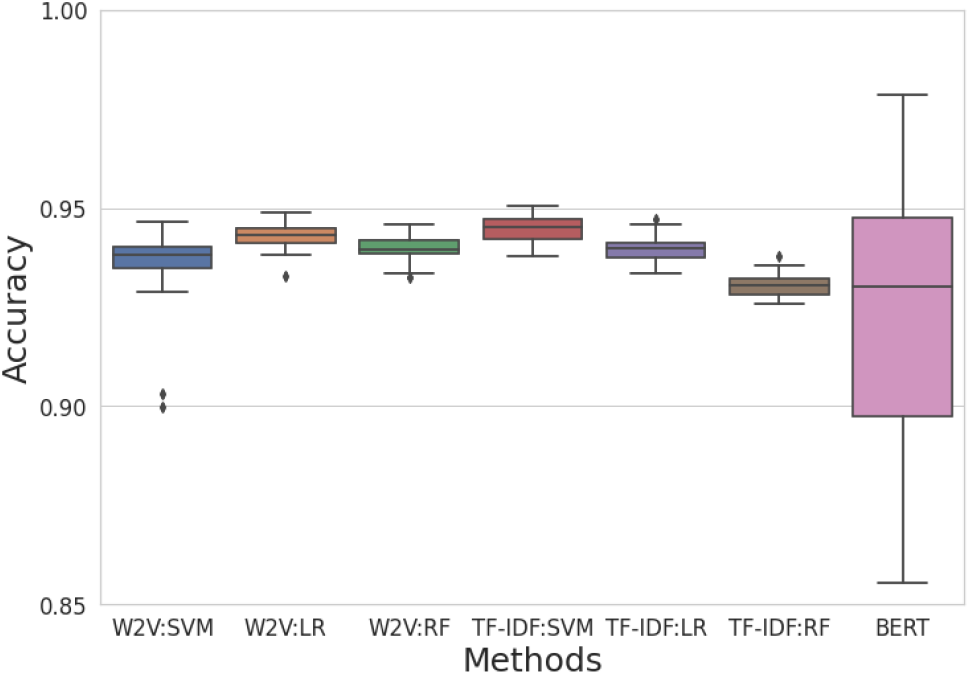
The accuracy of machine learning methods (SVM: support vector machine, LR: logistic regression, and RF: random forest) on test data using vectors derived from two vectorization techniques (W2V: Word2Vec and TF-IDF) in comparison with the BERT method.

**Table 1.** The accuracy, precision, recall, and F1-score of machine learning methods on test data using vectors derived from two vectorization techniques. The values in parentheses are the standard deviation. SVM: support vector machine, LR: logistic regression, RF: random forest.

| Vectorization | Methods | Accuracy (%) | Precision (%) | Recall (%) | F1-score (%) |
| --- | --- | --- | --- | --- | --- |
| Word2Vec | SVM | 93.6 (1.0) | 94.7 (1.9) | 92.5 (3.4) | 93.6 (1.2) |
|  | LR | 94.3 (0.3) | 94.6 (0.5) | 94.1 (0.5) | 94.4 (0.3) |
|  | RF | 94.0 (0.3) | 93.9 (0.5) | 94.2 (0.5) | 94.1 (0.3) |
| TF-IDF | SVM | 94.5 (0.4) | 94.8 (0.4) | 94.3 (0.5) | 94.5 (0.4) |
|  | LR | 94.0 (0.3) | 94.9 (0.5) | 93.1 (0.5) | 94.0 (0.3) |
|  | RF | 93.1 (0.3) | 94.1 (0.5) | 92.0 (0.5) | 93.0 (0.3) |
| Word2Vec & TF-IDF | SVM | **95.0 (0.3)** | 95.3 (0.4) | 94.7 (0.4) | 95.0 (0.3) |
|  | LR | 94.9 (0.3) | 95.1 (0.5) | 94.8 (0.4) | 94.9 (0.3) |
|  | RF | 94.1 (0.3) | 93.8 (0.5) | 94.6 (0.5) | 94.2 (0.3) |
|  | BERT | 91.7 (5.3) | 93.9 (5.6) | 92.5 (3.0) | 92.3 (3.2) |

### 2.2 External validation

A final linear SVM model was tested on independent datasets, resulting in accuracies of 92.5%, 96.3%, and 98.3%, and F1-scores of 93.5%, 86.1%, and 75.6% for three test sets (Table 2). The SVM model was tested on four independent validation sets, resulting in accuracies of 92.0%, 96.2%, 98.3%, and 93.1%, and F1-scores of 92.4%, 82.9%, 75.0%, and 93.3%.

**Table 2.**
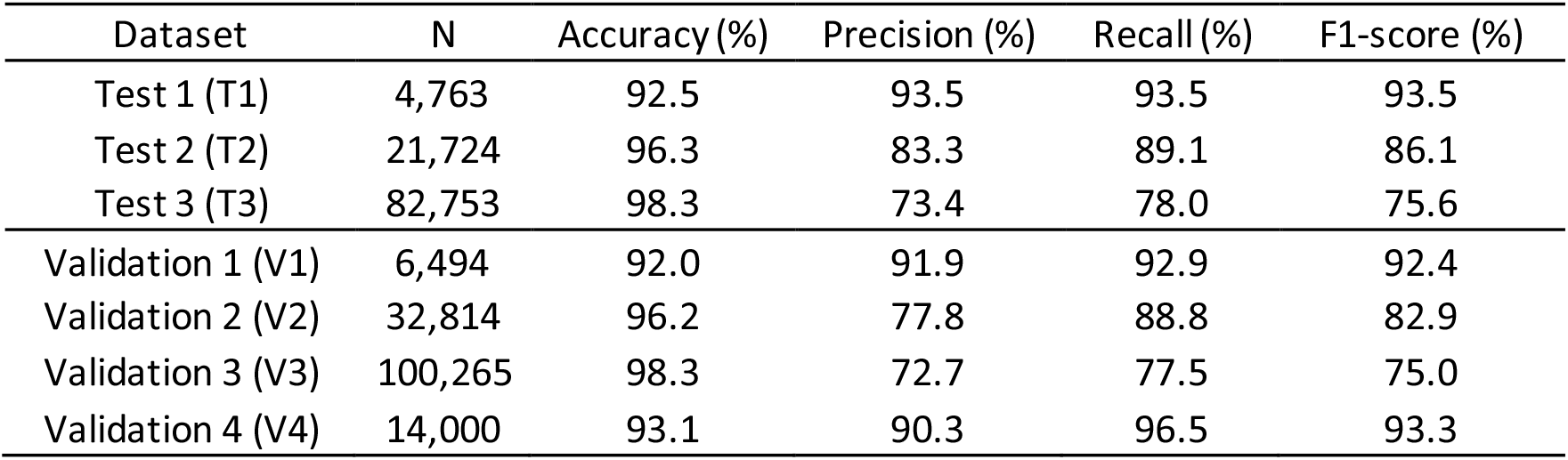
The accuracy, precision, recall, and F1-score of a final model assessed on three test and four validation datasets.

## 3. Conclusion

Machine learning methods using vectors derived from NLP text vectorization techniques were developed to classify literature related or unrelated to DILI. Machine learning models trained using the combined data of Word2Vec and TF-IDF vectors improved classification performance as compared to the models using Word2Vec or TF-IDF vectors alone. The developed analysis pipeline allows for easy adaptation to other NLP problems, facilitating the analysis of free-text documents by employing the integration of NLP techniques and machine learning.

